# Huge and variable diversity of episymbiotic CPR bacteria and DPANN archaea in groundwater ecosystems

**DOI:** 10.1101/2020.05.14.094862

**Authors:** Christine He, Ray Keren, Michael Whittaker, Ibrahim F. Farag, Jennifer Doudna, Jamie H. D. Cate, Jillian Banfield

## Abstract

Candidate Phyla Radiation (CPR) bacteria and DPANN archaea are uncultivated, small-celled symbionts often detected in groundwater. However, variations in CPR/DPANN organism abundance, distribution, taxonomic diversity, and degree/nature of host association with groundwater chemistry remain understudied. Here, we performed genome-resolved metagenomic characterization of one agriculturally-impacted and seven pristine groundwater microbial communities in California, recovering 746 dereplicated CPR and DPANN genomes. Our finding of up to 31% CPR bacteria and 4% DPANN archaea in the pristine sites, which serve as local sources of drinking water, may hold health relevance, given growing awareness of the presence of CPR/DPANN organisms in human microbiomes and their association with disease. There is little species-level genome overlap across groundwater sites, indicating that CPR and DPANN communities are highly differentiated according to host populations and physicochemical conditions. Cryo-TEM imaging and genomic analyses indicate that CPR growth may be stimulated by attachment to the surface of host cells, and identified CPR and DPANN lineages with particularly prevalent and/or resilient host cell attachment. These results establish the huge but site-specific diversity of CPR bacteria and DPANN archaea coexisting with diverse hosts in groundwater aquifers, and raise important questions about potential impacts on human health.

## Introduction

Recent metagenome-enabled phylogenomic analyses have led to the classification of two groups of organisms which lack pure culture representatives: the candidate phyla radiation (CPR) bacteria and DPANN archaea^1–4^. Although immense diversity exists within these two groups, CPR and DPANN organisms share conserved traits indicative of a symbiotic lifestyle: ultra-small cell sizes, small genomes, and minimal biosynthetic capabilities^5–9^. Episymbiosis (i.e. surface attachment) with prokaryotic hosts has been confirmed using co-cultures of Saccharibacteria with Actinobacteria^10^, Nanoarchaeota with Crenarchaeota^11,12^, and Nanohaloarchaeota and ‘ARMAN’ with Euryarchaeota^13–15^, though one case of endosymbiosis has been observed for a member of CPR superphylum Parcubacteria living within a protist^16^. CPR and DPANN organisms are ubiquitous across Earth’s environments and can be abundant in groundwater, where they are predicted to contribute to biogeochemical cycling^2,4,8,9,17–22^. CPR bacteria have been found to persist in drinking water no matter the treatment method^23–25^, posing the intriguing question of whether groundwater is a source of CPR^10,26–29^ and DPANN^30^ organisms detected in human microbiomes.

Despite their detection in groundwater, the variation in abundance and distribution of CPR and DPANN organisms across groundwater environments, their roles within groundwater ecosystems, and their relationships with host organisms are not well characterized. In general, subsurface environments are difficult to sample and poorly described compared to surface environments, despite harboring an estimated 90% of all bacterial biomass^31^. Many studies have likely underestimated CPR/DPANN organism abundance due to their ability to pass through 0.2 μm filters widely used to collect cells, as well as mismatch of “universal” primers to divergent or intron-containing 16S rRNA genes^2,11,13^. Most of the available near-complete CPR and DPANN genomes come from studies of only two aquifers^2,17,20^. A larger genome-resolved metagenomic study across groundwater environments is needed to place CPR and DPANN organisms (the latter rarely genomically characterized in groundwater studies) in the context of groundwater microbial ecosystems.

Here, we performed metagenomic characterization of eight groundwater sites in Northern California (seven pristine and one agriculturally-impacted, referred to as *Ag*). A total of 2,007 dereplicated genomes (≥70% complete and ≤10% contamination) were reconstituted, including 540 CPR and 206 DPANN genomes. This genome set enabled average nucleotide identity (ANI) based assessment of similarity between communities and metabolic profiling, shedding light on the influence of seasonal changes on metabolic capacities and the potential metabolic roles of CPR/DPANN organisms. Time series data from *Ag* showed a remarkably high abundance and diversity of CPR and DPANN organisms^32^ that were stably maintained over 15 months. Episymbiotic attachment of ultra-small cells to other microorganisms was observed using cryogenic transmission electron microscopy (cryo-TEM) imaging of *Ag* groundwater. Analyzing the distribution and replication rates of CPR genomes in different groundwater size fractions gave insight into the degree of episymbiotic attachment to hosts and correlation with CPR cell replication rates.

## Results

### Groundwater metagenome sampling and assembly

The planktonic fractions of eight groundwater communities in Northern California were sampled from 2017-2019 (Fig 1A) with bulk filtration (0.1 μm filter) and serial size filtration (2.5, 0.65, 0.2, and 0.1 μm filters). This enterprise required pumping 400-1200 L of groundwater from each site through a purpose-built sequential filtration apparatus to recover sufficient biomass for deep sequencing of each size fraction (Methods). One site (*Ag*) is an agriculturally-impacted, river sediment-hosted aquifer and the remaining sites are pristine groundwater aquifers hosted in a mixture of sedimentary and volcanic rocks (*Pr1* through *Pr7*, numbered in decreasing order of total CPR and DPANN organism abundance). Based upon the high abundance and diversity of CPR/DPANN organisms found at *Ag* in a previous metagenomics study^32^, we performed time series sampling over 15 months. All sites were sampled at shallow depths (<100 m).

Binning of bacterial and archaeal genomes from metagenomic data was performed using four different binning algorithms/techniques based on GC content, coverage, presence/copies of ribosomal proteins and single copy genes, tetranucleotide frequencies, and patterns of coverage across samples (see Methods). The highest quality bins were chosen using DASTool^33^ and then manually curated. All bins for a given site were dereplicated at 99% ANI^34^, resulting in a dereplicated set of 2,007 genomes across all sites (≥70% completeness and ≤10% contamination). The median genome completeness was >90% and up to 58% of metagenomic reads mapped to each site’s dereplicated genome set (see SI Table 1 and 2 for assembly and binning statistics). Of this dereplicated set, 540 and 206 genomes are classified as CPR bacteria and DPANN archaea, respectively (Supplementary Methods).

**Fig 1:**
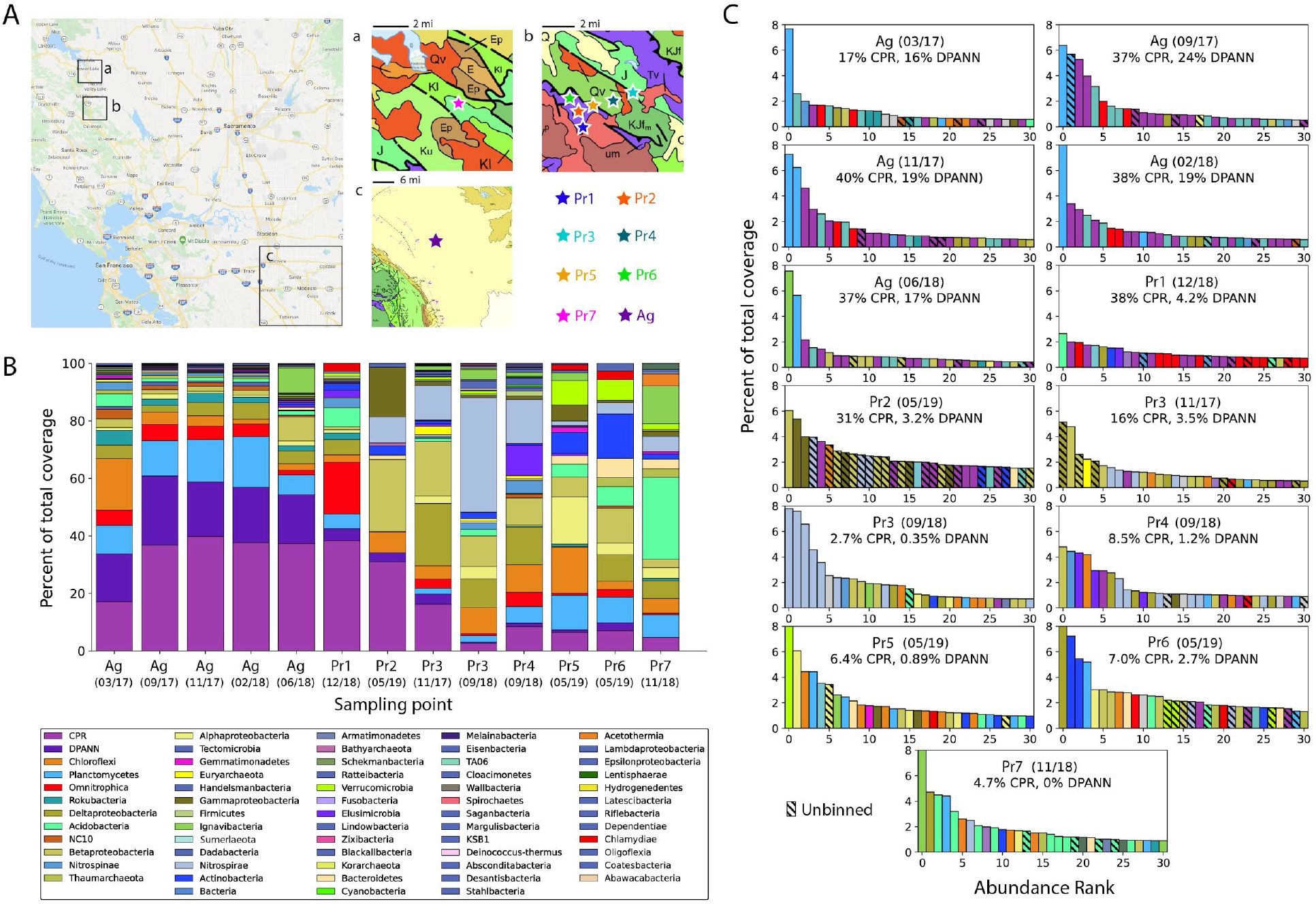
Sampling setup and rpS3 taxonomy-based overview of the community. **A,** Map of eight Northern California groundwater sites sampled in this study. Insets show geological maps of sampled areas. Qv = marine sedimentary and metasedimentary rocks (Cretaceous-Jurassic); J = marine sedimentary and metasedimentary rocks (Jurassic); Q = marine and nonmarine (continental) sedimentary rocks (Pleistocene-Holocene); KJfm = marine sedimentary and metasedimentary rocks (Cretaceous-Jurassic); um = plutonic (Mesozoic). **B**, Phylum-level breakdown (with the exception of CPR and DPANN superphyla) of rpS3 genes detected in each site. Sampling dates for each site are shown as (mm/yy). **C**, Rank abundance curves showing the 30 rpS3 genes with highest relative coverage identified for each site. Hatched bars indicate an unbinned rpS3 gene.

### Groundwater communities are highly distinct and vary dramatically in abundance and diversity of CPR/DPANN organisms

To evaluate how microbial community composition and diversity vary across groundwater sites, we used ribosomal protein uS3 (rpS3) as a single copy marker gene to broadly survey community composition^35^. Comparison of rpS3 genes against recovered genomes indicates that, with the exception of *Pr2*, the majority of the most abundant organisms in each site are represented by genome bins (Fig 1B, hatched bars indicate unbinned rpS3 genes). We find that all groundwater communities are distinct in phylum-level composition (Fig 1B and 1C), including in their populations of CPR and DPANN organisms (Fig 2A), though a few CPR and DPANN lineages are fairly ubiquitous across sites (SI Fig 1). Across all sites, CPR and DPANN organisms represent 3-40% and 0-24% of the communities, respectively. The abundance of DPANN archaea in *Ag* groundwater (10-24%) is significantly higher than in the pristine sites, where DPANN comprise <5% of the community. Across all sites, genomes were recovered from 58 of 73 currently identified phylum-level lineages within the CPR^4^ and from 6 out of 10 currently identified phylum-level lineages within the DPANN radiation (Fig 2B). In particular, recovered CPR genomes from *Ag* groundwater span most of the diversity within the CPR (Fig 2B, filled black circles). Based upon criteria for 16S rRNA gene sequence identity (<76% for phylum-level^19,36^) and concatenated ribosomal protein phylogenetic placement^2^, we defined two new phylum-level lineages within the CPR, each consisting of sequences from *Ag* and *Pr1* groundwater (SI Table 3). We propose the names Ca.

Genascibacteria and Ca. Montesolbacteria for these novel phylum-level lineages (highlighted in grey in Fig 2B), based upon the two sites where the representative sequences were found.

**Fig 2:**
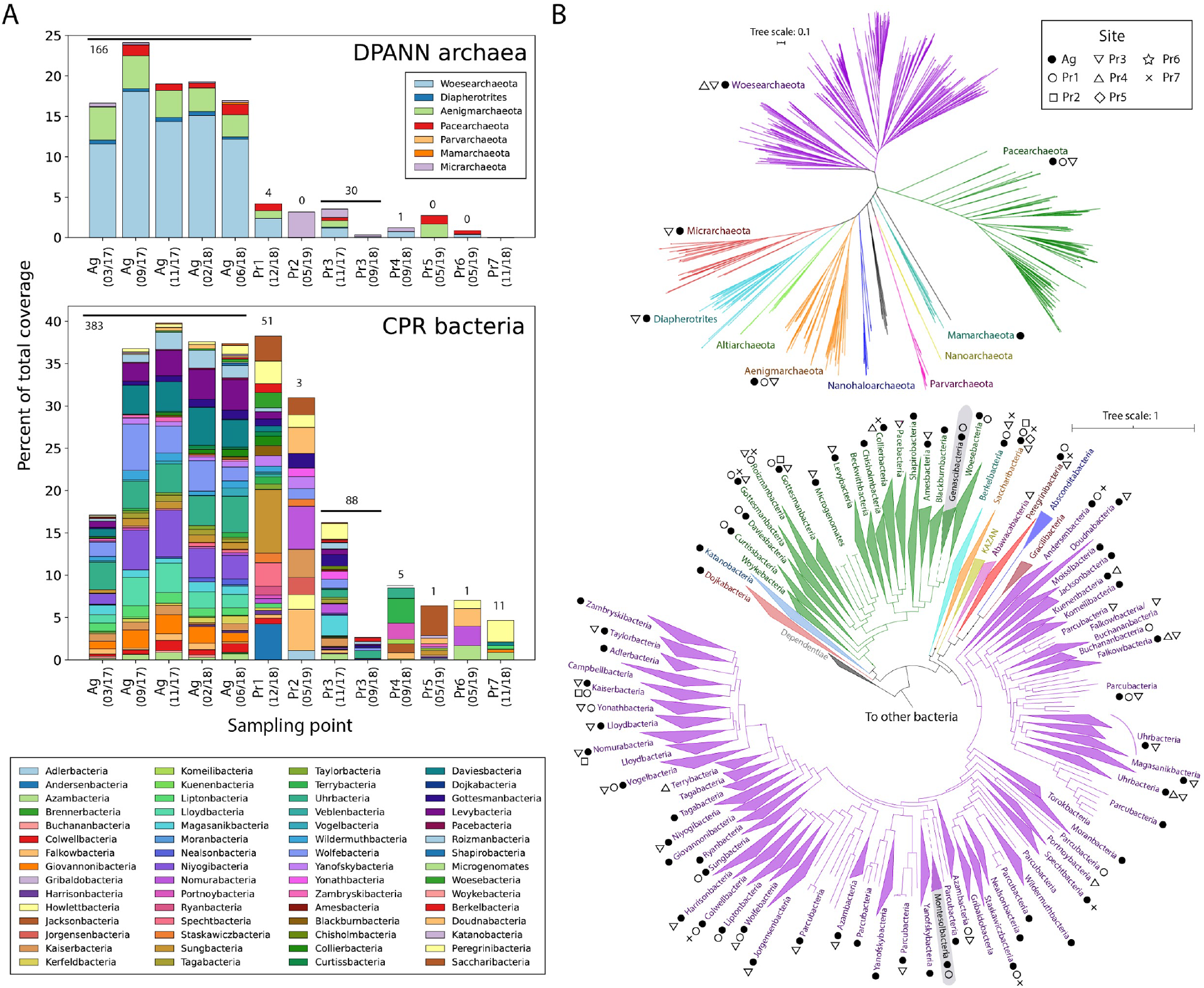
Diversity and distribution of CPR and DPANN organisms across groundwater sites. **A**, Abundances (rpS3-based) of phylum-level lineages within DPANN (above) and CPR (below) in all sites. Numbers above the bars indicate the number of high quality, dereplicated DPANN or CPR genomes recovered from metagenomic reads. **B**: Maximum likelihood phylogenetic tree of the DPANN radiation (top), based on 14 concatenated ribosomal proteins, and of the CPR (bottom), based on 15 concatenated ribosome proteins. Phylum-level lineages within the CPR (as previously defined^2^) are collapsed. Markers next to each lineage indicate the groundwater sites where at least one representative genome from that lineage was recovered. Novel CPR lineages Genascibacteria (within the Microgenomates superphylum in green) and Montesolbacteria (within the Parcubacteria superphylum in purple) are highlighted in grey.

Principal component analysis was performed on the relative coverage of phylum-level lineages, with scaling to mean of 0 and standard deviation of 1 (SI Fig 2). We see that *Ag* clusters separately from all pristine sites based upon strong and unique representation of organisms from Planctomycetes, Woesearchaeota, and several CPR lineages. *Pr1* is the site that clusters closest to *Ag* based on shared representation of Omnitrophica, while the remaining six pristine sites are differentiated by representation of Beta- and Deltaproteobacteria, Chloroflexi, and Nitrospirae. Change over time of the *Ag* groundwater community is discussed later, while differences between the two time points sampled for *Pr3* are discussed in Supplementary Note 1.

To assess groundwater community similarity at the genome level, we used ANI to cluster^34^ the 2,007 genomes from this study with 3,044 genomes from previous studies of two groundwater sites rich in CPR and DPANN organisms: Crystal Geyser, adjacent to the Green River, Utah^20,37,38^, and an aquifer adjacent to the Colorado River in Rifle, CO^2,17^. At the strain level (>99% ANI), there is very little similarity between genomes of analyzed sites, with most pairs of sites sharing one or no strains despite the fact that sites *Pr1* through *Pr6* are located in close proximity (~1 km between neighboring sites) and sites *Pr1, Pr2, Pr5*, and *Pr6* are all hosted in plutonic rock. The sole pair of sites that share more than a few strains (>99% ANI) is *Pr1* and *Pr7*, which share 44 strains, including 7 CPR bacterial strains (SI Fig 3). It is unlikely that the aquifers of these two sites are connected as they lie on separate sides of Putah Creek, a major hydrological feature. Additionally, we do not attribute this observed genome similarity to index hopping during sequencing (see Supplementary Note 2). Even at the species level (>95% ANI^39^), most pairs of analyzed sites share no more than one species in common (SI Fig 3). The overall lack of similarity between these ten groundwater sites--at the phylum level, species level, and strain level-- indicates a high degree of specialization based on local hydrogeochemical conditions.

### Potential roles of CPR and DPANN organisms in nitrogen and sulfur cycling within groundwater communities

Because most CPR and DPANN organisms are predicted to be symbionts, it is likely that their metabolic roles within a community vary with the metabolic capacities of their host organisms. To investigate this relationship, we profiled all recovered genomes against KEGG and custom protein HMM databases^40^ (Methods). By comparing the relative coverage of genomes encoding different metabolic capacities, we were able to metabolically profile whole communities (Fig 3A and SI Fig 4).

**Fig 3:**
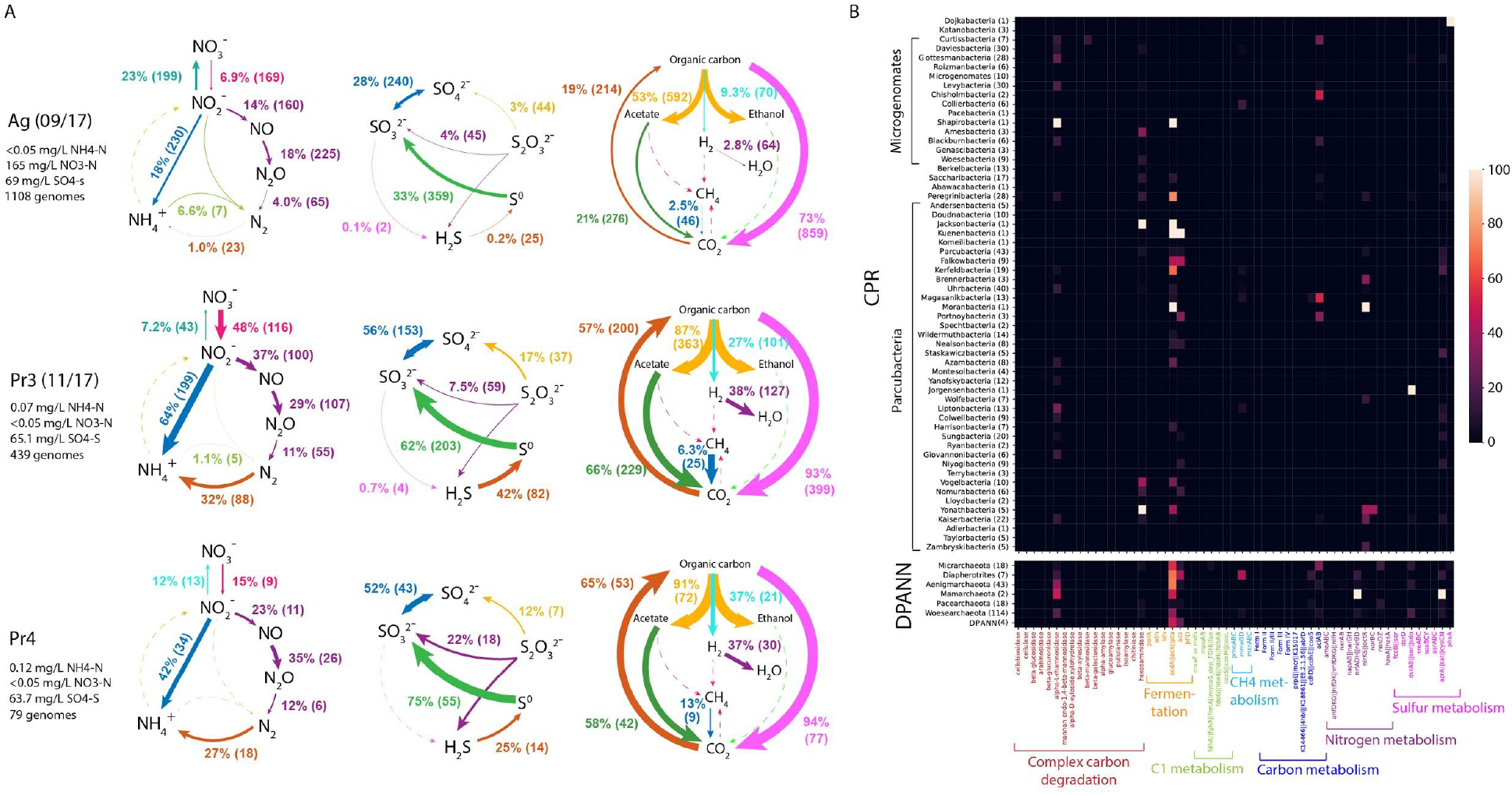
Metabolic profile of groundwater communities. **A,** Biogeochemical cycling diagrams profiling the community-level metabolism of five groundwater sites sampled in this study (all sites are shown in SI Fig 4). Listed next to each metabolic step are the total relative abundance of all genomes capable of carrying out the step, and the number of genomes containing the capacity for that step. Arrow sizes are drawn proportional to the total relative abundance of genomes capable of carrying out the metabolic step. **B,** Heatmap of 746 CPR and DPANN genomes from this study (rows are phylum-level lineages, and numbers in parentheses are the number of genomes recovered), showing the percentage of genomes containing key genes required for various metabolic and biosynthetic functions (columns).

Nitrogen input from agricultural waste and farming activities differentiates *Ag* from the pristine groundwater sites^32^ and as expected, *Ag* clusters separately from *Pr1-Pr7* in principal component analysis of community-level metabolic capacities (SI Fig 5). Ammonia is oxidized by seven Planctomycetes capable of anammox (comprising 8% of the community), resulting in low levels (<0.05 mg/L) of ammonia in *Ag* groundwater (Fig 3A). The *Ag* community encodes significantly greater capacity for nitrite oxidation than nitrate reduction (23% versus 7% of the community), consistent with measurements of high nitrate levels (165 mg/L NO3-N) and low nitrite levels (<0.05 mg/L NO2-N) (Fig 3A). Most groundwater communities sampled here (SI Fig 4) have incomplete capacity for denitrification, with far fewer genomes encoding the required genes for the final step of denitrification (nitrous oxide reduction) compared to previous steps. *Pr3*, *Pr4*, and *Pr6* are the only sites where >10% of the community can reduce nitrous oxide (Fig 3A). *Pr3* and *Pr4* also exhibit significantly higher capacity for nitrogen fixation, thiosulfate disproportionation, sulfide oxidation, and carbon fixation than the other groundwater communities investigated here (Fig 3A). While *Pr3* and *Pr4* have little species-level overlap, their similarity in community-level metabolic capacities may reflect their proximity (<1 km) and similar groundwater chemistry (levels of NH_4_-N, NO_3_-N, NO_2_-N, and SO_4_-S).

We further examined CPR and DPANN genomes in detail to assess what metabolic roles they may be playing in groundwater environments (Fig 3B). The presence of nitrite reductase *nirK* in 19 CPR genomes and 4 DPANN genomes as well as the presence of *nosD* in 11 DPANN genomes across sites suggest a complementary or accessory role of many CPR and DPANN lineages in different steps of denitrification (consistent with previous identification of *nirK* genes within Parcubacteria^19,41^). Additionally, 13 DPANN genomes in *Ag* groundwater encode for the small subunit of nitrite reductase (*nirD*) but lack the catalytic large subunit *nirB*, suggesting an accessory role of DPANN organisms in nitrite reduction to ammonia. In *Ag, Pr1*, and *Pr3*, we found that 30 DPANN genomes and 3 CPR genomes encode for sulfur dioxygenase *sdo*, while dozens of diverse CPR and DPANN genomes encode for *sat, cysC*, and *cysN* genes involved in sulfate reduction, suggesting a potential role of CPR and DPANN organisms in transformations to sulfite.

### Agriculturally-impacted dairy farm is a stable incubator of CPR bacteria and DPANN archaea

Time series and size filtration sampling of *Ag* groundwater allowed us to investigate the dynamics of a community rich in CPR and DPANN organisms (Fig 4A). Non-metric multidimensional scaling (NMDS) ordination (Methods) shows that, as expected, we see that most CPR and DPANN genomes cluster together and away from other bacteria and archaea, distinguished by prevalence in the 0.1-0.2 μm size fraction (Fig 4B). There is no significant clustering of genomes by sampling time in ordination space and the median root-mean-square deviation (RMSD) of genome relative abundances over time is ~0.002 (Fig 4C), indicating a very stable community at the level of genomes (dereplicated at 99% ANI). Inspection of individual genomes with an RMSD >0.004 (Fig 4D and 4E) showed that the most abundant organism, a Planctomycetes, peaked during 02/18. Several CPR bacteria peaked in abundance when the Planctomycetes abundance was low, suggesting a possible parasitic CPR-host relationship. Two Ignavibacteria and Betaproteobacteria organisms exhibited peak abundance on 06/18 (Fig 4D), while a similar trend is observed for an Uhrbacteria (Fig 4E), suggesting a potential commensal or mutualistic CPR-host relationship.

**Fig. 4:**
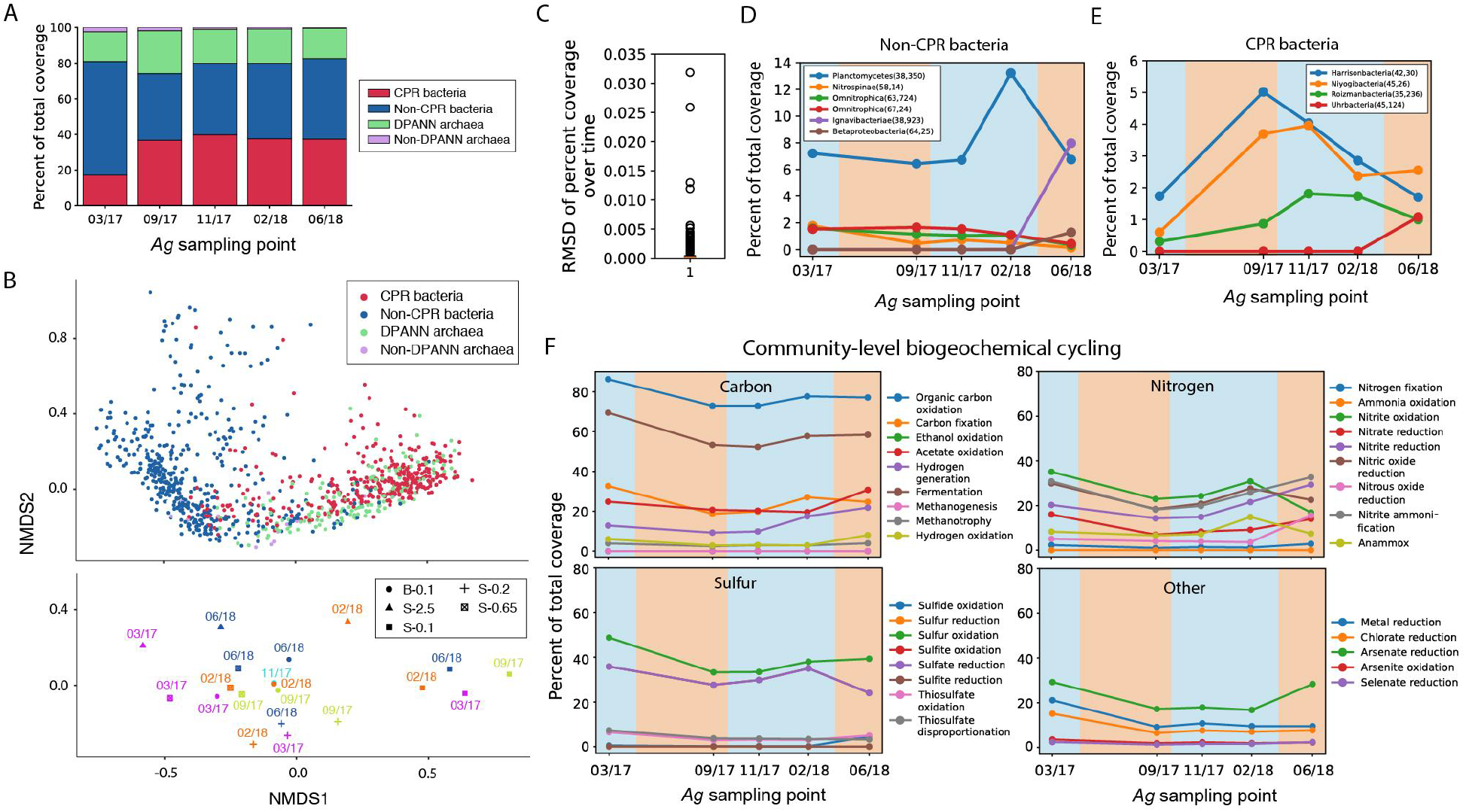
Genasci groundwater microbial community composition over time. **A**, Relative abundances of non-CPR bacteria, CPR bacteria, DPANN archaea, and non-DPANN archaea genomes in *Ag* over time. **B,** Non-metric multidimensional scaling (NMDS) analysis of *Ag* genome relative abundances in all size fractions over all time points. Position of genomes in ordination space are shown in the top graph, while the centroid of genomes belonging to a certain size fraction sampled at a certain date are shown in the bottom graph. **C,** Boxplot showing the root-mean-square deviation of relative abundance of all genomes in the bulk filter (whole community on a 0.1 μm filter) over time. The median RMSD (orange line) is <0.001, indicating that there is little variation in relative abundance over time for individual genomes in GD. **D,E:** Relative abundance over time for non-CPR bacteria (**D**) and CPR bacteria (**E**) that have an RMSD > 0.004. Genomes are identified in the legend by phylum, percent GC, and coverage in the original time point the representative genome was derived from (the latter two in parentheses). **F,** Variation in *Ag* community-level capacity (total relative abundance of all genomes capable of a broad metabolic function) for carbon, nitrogen, sulfur, and miscellaneous element cycling over time. Panels **D, E, F:** blue and orange backgrounds indicate the rainy and dry seasons in Northern California.

Examination of changes in metabolic cycling capacities in *Ag* groundwater over time indicate higher community capacity (5-10% relative abundance) for organic carbon oxidation, carbon fixation, fermentation, nitrite oxidation, nitric oxide reduction, and sulfate reduction during the rainy season (Fig 4F, blue background) compared to the dry season (Fig 4F, orange background). Furthermore, we see a higher increase in these metabolic capacities during the 2016-17 compared to the 2017-18 rainy season (Fig 4F), which may reflect a major difference in rainfall (17.93” in 2016-17 versus 7.87” in 2017-18)^42^. Overall, we find that *Ag* groundwater is a remarkably stable incubator for high abundance and diversity of both CPR and DPANN organisms, but the microbial community is not stagnant in its metabolic capacities, which vary between rainy and dry seasons.

### Cryo-TEM imaging shows pili-mediated episymbiotic interactions between ultra-small cells and host cells

To observe CPR/DPANN-host interactions in the *Ag* groundwater community with microscopy, we avoided cell concentration with dead end filtration, due to potential induction of artifactual associations between cells caught on a filter. Instead, we used tangential flow filtration (TFF) to concentrate cells from groundwater and preserved them by cryo-plunging them in liquid ethane on-site for later characterization by cryogenic transmission electron microscopy (cryo-TEM) (Methods).

Many ultra-small cells (longest dimension <500 nm) were observed attached to the surface of larger host cells (Fig 5A). The ultra-small cells have cell envelopes decorated by pili (Fig 5, white arrows), some of which extend into the corresponding host cell, potentially mediating episymbiont-host interaction (Fig 5, white dashed boxes). The presence of pili identifies these ultra-small cells as CPR bacteria, since components of type IV pili systems are present in the majority of CPR bacterial genomes and missing in DPANN genomes^4,43^. At the CPR/host contact region in Fig 5E and H, the host cell envelope appears to be thickened, while the CPR bacterial cell envelope is thinned. For multiple CPR/host pairs, a line of higher density is observed at the cell interface (Fig 5D, E, and I, orange arrows), similar to what has previously been observed at tight interfaces between archaeal ARMAN (DPANN) cells and their *Thermoplasmatales* hosts^44^. The host in Fig 5A has multiple CPR bacterial cells directly attached to its cell envelope which appear to be in the process of dividing (Fig 5B, 5E, 5F), raising the possibility that CPR replication is correlated with host attachment (discussed in the next section). Other ultra-small cells were observed to be associated with/in close proximity to lysed cells, suggesting that some CPR/DPANN organisms scavenge resources from dead cells or act parasitically^10^ (SI Fig 6). Overall, cryo-TEM imaging of TFF-concentrated groundwater shows that some CPR bacteria in *Ag* groundwater are episymbionts of prokaryotic hosts, attaching through pili-like structures.

**Fig 5:**
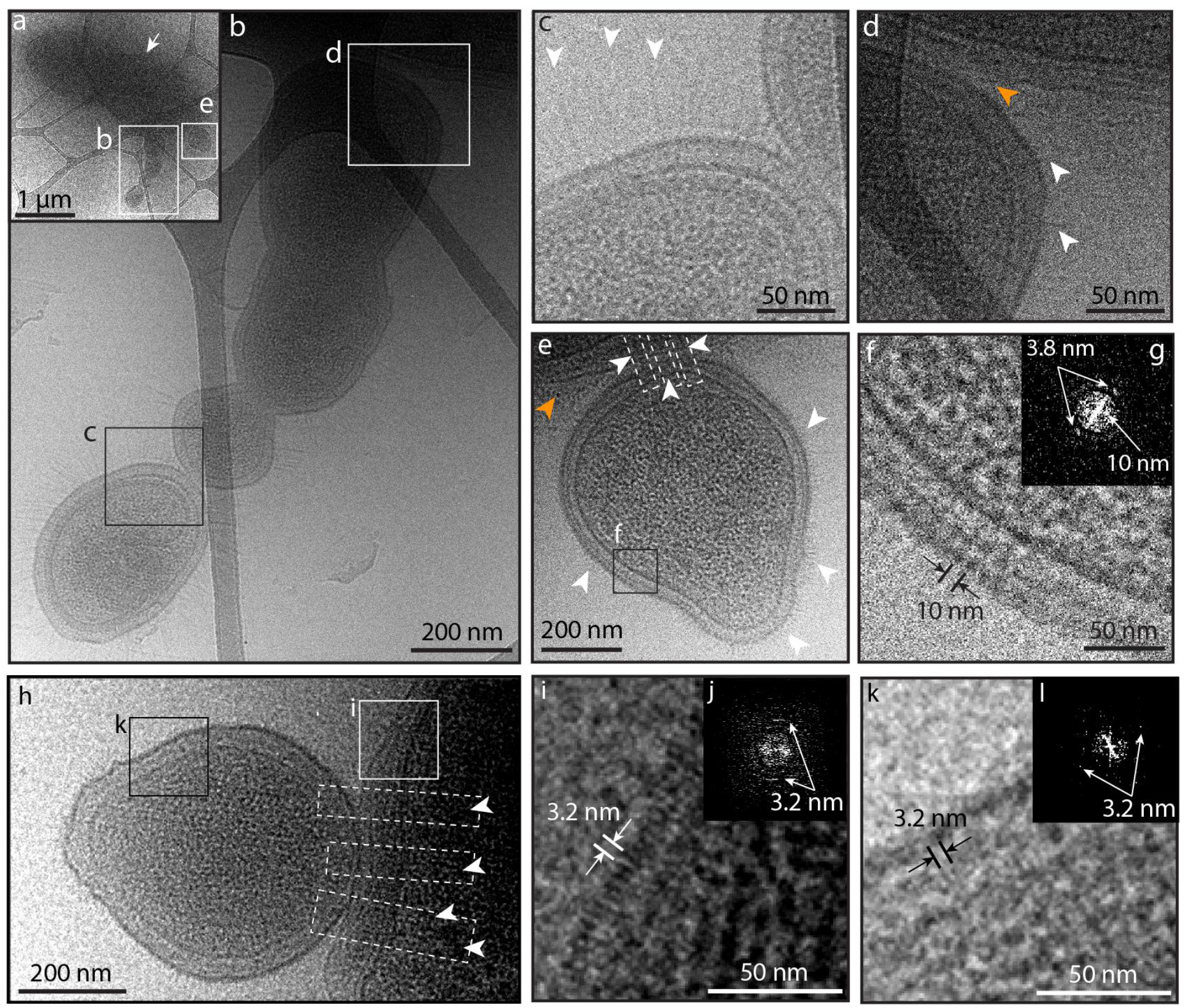
Cryo-TEM images of Genasci groundwater cells concentrated with tangential flow filtration. **A:** Image of larger host cell (white arrow) with multiple ultra-small cells (panels b and e) attached. **B:** Inset of panel a showing a chain of ultra-small cells attached to the surface of the larger host cell. **C:** Inset of panel b showing the contact region between two ultra-small cells and the pili-like appendages decorating their surfaces (white arrows). **D:** Inset of panel b showing the contact region between an ultrasmall cell and host cell. Pili-like appendages are indicated with white arrows. **E:** Inset of panel a showing a single ultra-small cell decorated by pili-like appendages (white arrows) attached to a host cell. Attachment may be mediated by pili-like appendages that extend from the ultra-small cell into the host cell (white boxes). **F:** Inset of the membrane from the ultra-small cell in panel e, with the membrane structure showing a clear periodicity measured to be 10 nm as well as a periodicity of 3.8 nm evident in Fourier space (panel **G**). **H:** Image of a host cell with a single ultra-small cell attached. Several pili-like appendages extend from the ultra-small cell into the host cell (white boxes and arrows). **I:** Inset of panel h showing the host cell envelope, the outer layer of which exhibits a periodicity of 3.2 nm (Fourier transform in panel **J**). **K**: Inset of panel h showing the ultra-small cell envelope, which also exhibits a periodicity of 3.2 nm in the outer layer (Fourier transform in panel **l**). In all images, orange arrows indicate lines of high density observed at the contact interface.

An important question regarding the biology of CPR bacteria relates to the nature of their cell envelope and the degree to which it resembles that of host cells to which they attach. Genomic analysis indicates that CPR bacteria cannot *de novo* synthesize fatty acids^4^ but do possess fatty acid-based membrane lipids^45^, raising the possibility that CPR bacteria symbiotically receive lipids or lipid building blocks from host organisms. The host-attached CPR bacterium in Figure 5E has a surface layer (S-layer) with a periodicity of 3.8 and 10 nm, but lacks an outer membrane expected for Gram-negative bacteria (Fig 5F and 5G), consistent with previous cryo-TEM images of groundwater CPR bacteria^43^. Meanwhile, the host’s cell envelope appears to have two lipid layers, suggesting a Gram-negative structure (Fig 5D and E). In Figure 5H, from a different host-CPR pair, we also resolve two lipid layers in the host cell envelope and no outer membrane in the CPR bacterium. Interestingly, in this case, a periodicity of 3.2 nm is detected in the outermost layers of both the host cell (Fig 5I) and the attached CPR bacteria (FIg 5K), but it is unclear whether the two cells’ outer layers have the same structure. Based on an apparent lack of an outer membrane, CPR bacteria in *Ag* groundwater have cell envelopes that do not resemble those of Gram-negative bacteria, but seemingly can attach to Gram-negative hosts.

### Size fractionation gives insights into attachment between CPR/DPANN and host cells

We analyzed how CPR/DPANN organisms were distributed among size fractions, which should reflect two factors: cell size and attachment to larger host cells. Most microbes outside the CPR and DPANN groups are present in the 0.65-2.5 or 2.5+ μm size fractions (Fig 4B). Due to their small cell size, a CPR/DPANN cell present in the 2.5+ μm size fraction is likely to be attached to a larger organism in the same size fraction, while a CPR/DPANN cell present in the 0.1-0.2 μm size fraction is likely to be unattached. While pumping groundwater through filters may detach CPR/DPANN cells from hosts, substantial coverage of CPR/DPANN organisms in the 2.5+ μm and 0.65-2.5 μm size fractions indicate that a fair number of CPR/DPANN cells remain host-attached through filtration. Additionally, we consider it likely that pili penetrating from CPR cells into the host (Fig 5) are strong enough to resist disruption. We therefore consider the distribution of CPR/DPANN organisms among size fractions as indicative of host attachment, i.e. what fraction of CPR/DPANN cells are host-associated and the resistance of the cell-cell association to the disruptive effects of filtration.

To assess the distribution of organisms among size fractions, the absolute number of cells represented by a genome was estimated from the genome relative abundance and the mass of DNA extracted from each size fraction (Methods). As expected, CPR and DPANN lineages had significantly higher cell counts in the 0.1-0.2 μm fraction compared to the 2.5+ μm fraction, while other lineages exhibited the reverse trend (paired t-test, Fig 6A). Two notable exceptions are CPR lineage Kerfeldbacteria and DPANN lineage Pacearchaeota, which were enriched on the 2.5+ μm size fraction relative to the 0.1-0.2 μm size fraction (paired t test, p = 0.0048 and 0.014) (Fig 6A), indicating that a high fraction of these populations is host-attached and/or the attachment is more resistant to the disruptive effects of filtration compared to other CPR/DPANN lineages. Pacearchaeota genomes encode especially minimal metabolic capacities among DPANN lineages (Fig 3B), suggesting a heavy dependence upon host resources^46^. An additional CPR lineage, Woesebacteria, was found to have significantly higher cell counts in the 2.5+ μm size fraction versus the 0.2-0.65 μm size fraction (SI Fig 7).

**Fig 6:**
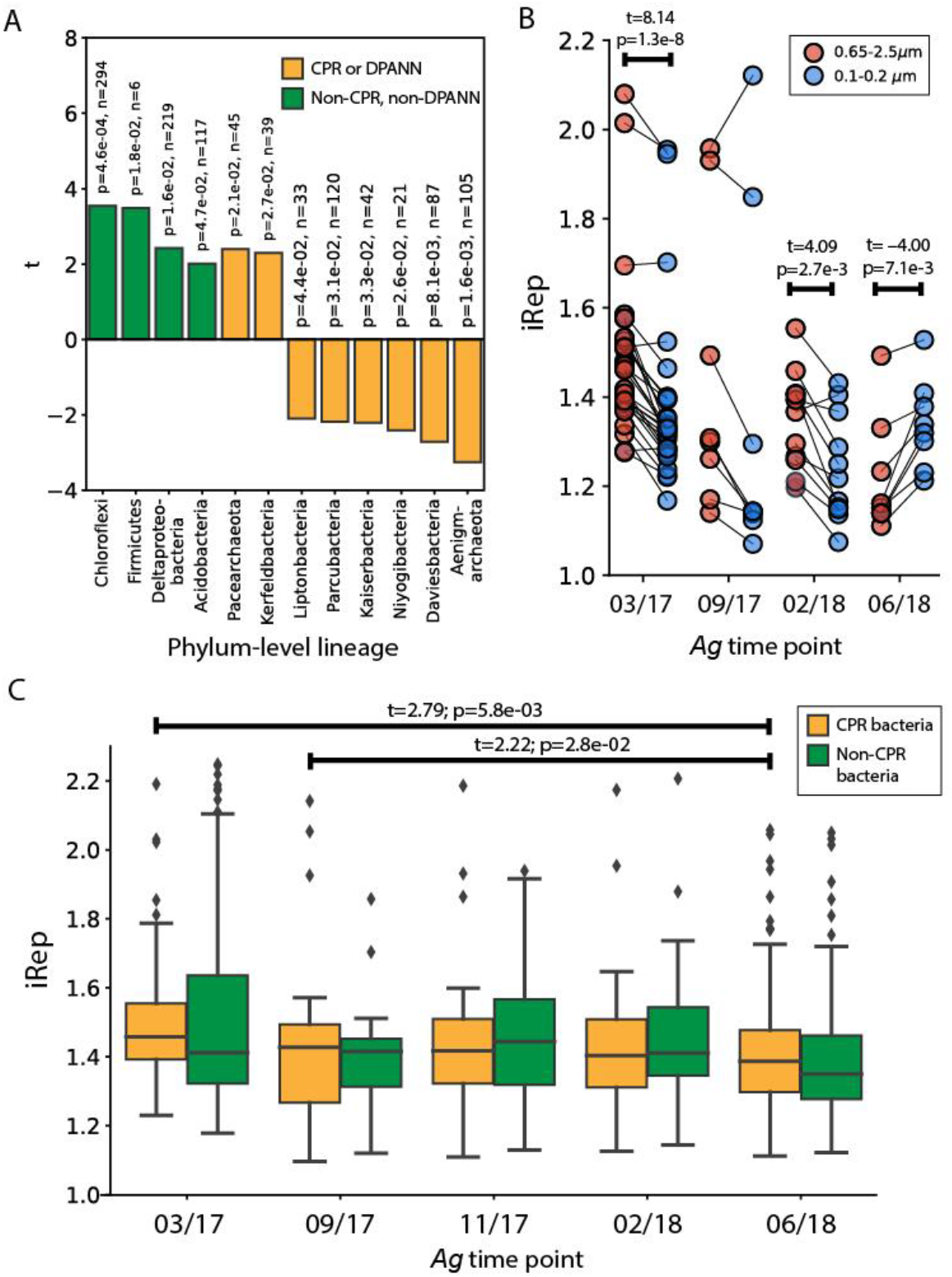
Analysis of CPR-host attachment in Genasci groundwater. **A**, Results from a paired t-test on estimated cell counts of genomes in the largest (2.5+ μm) and smallest (0.1-0.2 μm) size fractions after serial size filtration. A positive t statistic indicates enrichment on the 2.5+ μm compared to the 0.1-0.2 μm size fraction. Values listed by each bar are the calculated p value and sample size (n) for each phylumlevel lineage. **B,** Calculated iRep values for CPR bacteria genomes in the 0.65-2.5 μm size fraction versus the 0.1-0.2 μm size fraction, across all *Ag* sampling points. The statistically significant results (p < 0.05) of a paired t test on iRep values between the two size fractions are shown above the box plots. **C**, Calculated iRep values for bacteria caught in the bulk 0.1 μm filter (whole community filtration). The statistically significant results (p < 0.05) of an independent t test on iRep values of CPR bacteria between all possible pairs of sampling points are shown above the box plots.

As noted above, many of the attached ultra-small cells appear to be dividing (Fig 5A, panel b), suggesting that attachment to a host and acquisition of metabolic resources may stimulate division of CPR/DPANN organisms. To investigate this hypothesis, we calculated instantaneous replication rates (iRep values^47^) for CPR genomes in the *Ag* groundwater bulk community and size fractions. Replication in archaea does not generally proceed in a bidirectional manner, so archaeal genomes were excluded from iRep analysis. Tracking iRep values of CPR genomes across size fractions, we see that at two sampling points--03/17 and 02/18, the height of the rainy season--CPR organisms exhibit higher replication rates in the 0.65-2.5 μm fraction than the 0.1-0.2 μm size fraction (Fig 6B). During the height of the 2016-17 rainy season (03/17) and the end of the dry season/beginning of the next rainy season (09/18), we found that CPR bacteria as a whole exhibited higher replication rates than in the height of the 2018 dry season (06/18) (Fig 6C). Significant differences in replication rates were not observed between any point and the height of the rainy season on 02/18 (Fig 6C), which may be explained by much higher rainfall in the 2016-17 rainy season (17.93”) than in the 2017-18 rainy season (7.87”)^42^. Together, these findings support the deduction that CPR cell replication is stimulated by host attachment, and indicate that this may be more prevalent during the rainy season compared to the dry season.

## Discussion

In this study, we sampled one agriculturally-impacted and seven pristine groundwater sites in Northern California, hosted in a range of rock types and sourced from multiple aquifers. The recovery of 746 draft quality CPR and DPANN genomes was enabled by filtering groundwater onto 0.1 μm filters as well as sequentially onto a series of 2.5, 0.65, 0.2, and 0.1 μm filters. Large volumes of groundwater (400-1200 L per site) were pumped in order to collect sufficient biomass for deep sequencing of all filters. Different size fractions provided genomic resolution of different community members and provided differential coverage patterns for constraining bin assignment. The recovered genomes derive from most of the major lineages within the CPR and DPANN radiations and from two apparently new phylum-level lineages within the CPR, named Ca. Genascibacteria and Ca. Montesolbacteria. To our knowledge, only two previous studies have recovered and compared CPR bacterial genomes across multiple groundwater sites, and neither has looked at DPANN genomes. The number of draft quality CPR genomes in this study (540) far exceeds the number reported in these previous groundwater site comparison studies (71^19^ and 6^48^).

There is very little species-level overlap (defined as >95% ANI) between genomes recovered from this study or with genomes from previous studies of Crystal Geyser^20,38^ and Rifle^2,17^. The sole exception is a pair of sites, *Pr1* and *Pr7*, that share 44 species (>95% ANI) despite low potential for connection between the aquifers. The high level of differentiation between groundwater communities likely reflects species adaptation to different physiochemical conditions and suggets that further sequencing of groundwater sites is likely to reveal unexplored diversity within the CPR and DPANN radiations.

The pristine sites sampled here serve as sources of local drinking water. Notably, at the time of sampling, *Pr2* (Rattlesnake Spring), a popular source of public drinking water for over a century, contained over 30% CPR bacteria and 3% DPANN archaea. Currently, there is limited information about the presence and role of CPR and DPANN organisms in human and animal microbiomes. Detection of CPR lineage Saccharibacteria in multiple human body sites has been correlated with inflammatory bowel disease^26^, vaginosis^49^, and periodontitis ^50–52^, while CPR lineages Absconditabacteria and Gracilibacteria have been detected in human saliva and correlated with herpes viral titers^29,53^. Members of CPR superphylum Parcubacteria have been detected in the human gut^54^ and DPANN archaea have been detected in lung fluids^30^. The findings of the current study raise the possibility of drinking water as a source of CPR bacteria in human microbiomes, as CPR bacteria have been found to persist in drinking water after treatment^23–25^.

An important aspect of this study was the use of serial size filtration for fractionating CPR and DPANN cells according to host attachment. This allowed us to identify two lineages, Kerfeldbacteria in the CPR and Pacearchaeota within DPANN, with likely strong physical attachment to hosts. No genomic features identified Kerfeldbacteria as particularly distinct from other CPR, but Pacearcheaota genomes in this study and across environments have notably minimal metabolic capacity^46^. Size fraction data combined with cryo-TEM imaging of *Ag* groundwater also enabled us to associate higher CPR instantaneous replication rates with physical attachment to a host. We conclude that CPR organisms tend to grow and replicate after stimulation by host attachment and the availability of host-supplied resources.

The *Ag* groundwater microbial community, which includes organisms from most CPR and DPANN lineages, is remarkably stable at the strain level (>99% ANI), perhaps due to consistent, heavy input of carbon and nitrogen from agricultural waste. Though the *Ag* community composition is fairly stable, increases in community metabolic capacities and CPR bacterial replication rates occur during rainy seasons. Several factors may contribute to these changes during the rainy season: the onset of more anaerobic conditions in the groundwater, greater runoff from agricultural waste piles, increased volume of and changes in microbial composition of the cow manure after calves are born in the spring, and soil changes associated with the adjacent corn field that supplies much of the recharge to the sampled *Ag* well^32^. Analysis of abundance patterns over time identified potential parasitic as well as commensal/mutualistic relationships between several CPR lineages and Planctomycetes, Ignavibacteria, and Betaproteobacteria hosts. These observations provide a starting point for targeted cultivation of CPR and DPANN organisms based on conditions favorable to growth of putative hosts.

## Methods

### Groundwater sampling, chemistry measurements, and surface geology determination

Groundwater was pumped from each well using a submersible pump (Geotech Environmental Equipment) into a sterile container, and then pumped with a peristaltic pump into an apparatus custom built for filtering high volumes of water (Harrington Industrial Plastics) at a rate of 3.8-7.6 L/min. Prior to filtration, at least 100 L of water were pumped to purge the well volume and to flush the system. Polyethersulfone membrane filter cartridges designed for high volume filtration (Graver Technologies) from the ZTEC G series (0.1 μm and 0.2 μm), ZTEC B series (0.65 μm), and PMA series (2.5 μm) were used. When a sufficient volume of water had been filtered (400 L for bulk filtration and an additional 800 L for serial size filtration), filters were removed and stored on dry ice. Filters were stored in a −80 C freezer until processed. The surface geology of each sampling site was determined from the California Department of Conservation’s 2010 geologic map of California (https://maps.conservation.ca.gov/cgs/gmc/), rock fragments recovered during drilling (*Pr4*), and by on site geological surveys. Pumped groundwater was shipped on dry ice to the UC Davis Analytical Laboratory for water chemistry measurements of: EC, SAR, TOC, DOC, NH_4_-N, NO_3_-N, SO_4_-S (soluble S), HCO_3_, CO_3_, soluble Zn, Cu, Mn, Fe, Cd, Cr, Pb, Ni, K, Ca, Mg, Na, Cl, and B. Water chemistry measurements are shown in SI Table 4.

### DNA extraction and sequencing

The plastic housing was removed from the filter cartridges under sterile conditions and the filters were retained for DNA extraction. To extract DNA, either a quarter or a half of a filter were placed in Powerbead solution from the Qiagen DNAeasy PowerSoil kit (no bead beating was performed), then vortexed for 10 minutes with massaging to remove cells from the entire filter surface. After vortexing, the filter was removed, solution C1 (Qiagen DNAeasy PowerSoil Kit) was added to the Powerbead solution, and the solution was placed in a 65 ^o^C water bath for 30 minutes. The remainder of DNA extraction proceeded according to the Qiagen DNeasy PowerSoil kit manufacturer instructions, beginning with the addition of solution C2. Ethanol precipitation was performed to concentrate and purify the extracted DNA before sequencing. Genomic DNA was quantified using Qubit dsDNA High Sensitivity assay and when quantity permitted, DNA quality was assessed using agarose gel electrophoresis. Library preparation and sequencing were performed at the California Institute for Quantitative Biosciences’ (QB3) genomics facility and the Chan Zuckerberg Biohub’s sequencing facility. Libraries were prepared with target insert sizes of 400-600 bp. Samples were sequenced using 150 bp paired-end reads on either a HiSeq 4000 platform or a NovaSeq 6000 platform, with a read depth of ~10 Gbp per sample except for *Ag* 03/17 samples, which were sequenced at 150 Gbp.

### Metagenomic assembly

BBTools was used to remove Illumina adapters as well as PhiX and other Illumina trace contaminants^55^. Reads were trimmed using Sickle^56^ (version 1.33). Assembly was performed using MEGAHIT (version 1.2.9) with default parameters^57^, and the scaffolding function from assembler IDBA-UD^58^ was used to scaffold contigs. Scaffold coverage values were calculated using read mapping with bowtie2^59^. Only scaffolds >1 kb in length were considered for gene prediction and genome binning. Gene prediction was performed using Prodigal (version 2.6.3) with the ‘meta’ option^60^ and genes were annotated using usearch^61^ against KEGG^62,63^, Uniref100^64^, and UniProt^65^ databases. 16S rRNA genes were identified using a custom HMM^2^ and insertions of 10 bp or greater were removed. Prediction of tRNA genes was performed with tRNAscan-SE^66^.

### Genome binning, curation, and dereplication

Scaffolds were binned on the basis of GC content, coverage, presence/copies of ribosomal proteins and single copy genes, tetranucleotide frequency, and patterns of coverage across samples. A combination of manual binning on ggKbase (https://ggkbase.berkeley.edu/) and automated binning using CONCOCT^67^, Maxbin2^68^, and Abawaca2 were used to generate candidate bins for each sample. Best bins were determined using DASTool^33^ and manually checked using ggKbase to remove incorrectly assigned scaffolds according to the criteria listed above. Genomes were then filtered for completeness (>70%) using a set of 43 single copy genes previously used for the CPR^2,17^ and 48 single copy genes for DPANN, and contamination using checkM^69^ (<10%) (SI Table 2). The program dRep^34^ was used to dereplicate genomes from each site at 99% ANI (strain level), resulting in a representative set of 2,007 genomes across all sites. The median estimated genome completeness of each site’s representative set is over 90%, with 18-58% of each site’s raw reads mapping back to the representative set (SI Table 1).

### Genome and community-level metabolic predictions

To analyze the metabolic capacity of the sampled groundwater communities at both the genome and community level, the program METABOLIC^40^ (version 1.3) was used to search predicted ORFs against KEGG, TIGRfam, Pfam, and custom HMM profiles, to determine the presence/absence of metabolic capacities (based on KEGG modules) encoded by each genome, and to calculate the number and relative abundance of genomes in the community which encode for a metabolic capacity. Because metagenome-assembled genomes often encode for incomplete metabolic pathways, a given KEGG module was considered present within a genome if genes were identified for >75% of the reactions in the module. The presence/absence of critical marker genes (i.e. RuBisCO Forms I through IV for the CBB pathway) were also examined to help assess whether a particular KEGG module was encoded by a genome.

### Cryo-TEM sample preparation in the field

Cryo-TEM samples were prepared onsite at the *Ag* dairy farm on Feb 5, 2018. Approximately 30 L of pumped *Ag* groundwater was concentrated to a final volume of ~5 mL, using tangential flow filtration (TFF) (Millipore Pellicon Cassette Standard Acrylic Holder) with a 30 kDa ultrafiltration cassette (Millipore Pellicon 2 Biomax). Aliquots of 5 μL were taken directly from the suspensions and deposited onto 300 mesh lacey carbon coated Cu-grids (Ted Pella, #01895) that had been treated by glow discharge within 24 hours. Grids were blotted with filter paper and plunged into liquid ethane held at liquid nitrogen temperatures using a portable, custom-built cryo-plunging device^70^. Plunged grids were stored in liquid nitrogen prior to transfer to the microscope and maintained at 80 K during acquisition of all data sets.

### Cryo-TEM imaging

Imaging was performed on a JEOL–3100-FFC electron microscope (JEOL Ltd., Akishima, Tokyo, Japan) equipped with a FEG electron source operating at 300 kV. An Omega energy filter (JEOL) attenuated electrons with energy losses that exceeded 30 eV of the zero-loss peak prior to detection by a Gatan K2 Summit direct electron detector. Dose-fractionated images were acquired with a pixel size of 3.41 Å/pixel using a dose of 7.27 e^-^/Å^2^ per frame. Up to 30 frames per image were aligned and averaged using IMOD^71^ and image contrast was adjusted in ImageJ (v2.0.0).

### Analysis of cell distribution across serial size filters

To analyze distribution of *Ag* genomes across size fractions, we estimated total cell counts from genome relative coverage values and the total mass of DNA in each size fraction. The coverage of a genome is proportional to cell counts, so we assumed that a genome’s relative coverage is the fraction of total cells in the community represented by that genome. Therefore, the cell count of a genome was calculated as the product of the genome relative coverage and estimated total cells in the community.

Relative coverage of a genome in a given size fraction was calculated by mapping reads from the appropriate serial size filter (i.e. the 0.1 μm filter for the 0.1-0.2 μm size fraction) using bowtie2^59^. The total mass of DNA in each size fraction was measured by a Qubit dsDNA HS Assay on DNA extracted from each filter. From these two measurements, the total cell count of a genome in a given size fraction was calculated as *c* = *x* * *I* * *m*, where c = total cell count of a genome; x = relative coverage of a genome; l = cell counts per ng of DNA in the community; and m = ng of DNA extracted from the size fraction. The parameter l was estimated from the total set of representative genomes recovered from the environment, with the molecular weight of each genome calculated as number of base pairs * 650 Da/base pair.

To find significant differences in cell counts between two given size fractions (i.e. 2.5+ μm versus 0.1-0.2 μm), a paired t-test was performed on cell counts from each phylum with more than 5 representative genomes and with cell count distributions in each size fraction that did not deviate significantly from normality (assessed by plotting cell count distributions and performing a Shapiro-Wilks test).

### iRep analysis

Instantaneous replication rates were calculated for *Ag* bacterial genomes using iRep^47^ (version 1.1.14) with a tolerance of 3 mismatches per read. Reads from each size fraction were mapped to the bacterial genomes using bowtie2^59^.

## Data Availability

SRA accession numbers for metagenome reads are in SI Table xx. All metagenome-assembled genomes from this study are deposited in NCBI under Bioproject xx. Prior to access via NCBI, the data are available at: http://ggkbase.berkeley.edu/all_nc_groundwater_genomes (please note that it is necessary to register for an account by provision of an email address prior to download).

## Acknowledgements

We thank Edwin Genasci, Alan Book, Kurt Kritikos, Kendall and Dustin Fults, Peregrine Smith, and Justin Smith for access to their groundwater wells. We thank Cindy J. Castelle, Luis Valentin, Basem Al-Shayeb, Katherine Lane, Matthew Olm, Nikolin Oberleitner, Raphael Méheust, Alexander Jaffe, and Jacob West-Roberts for their assistance with sampling. We thank Cindy J. Castelle for assistance with archaeal phylogenetic analysis and Daniel Toso for help with collection of cryo-TEM data. C.H. was supported by the Camille and Henry Dreyfus Foundation Postdoctoral Fellowship in Environmental Chemistry. Funding for groundwater sampling and sequencing was provided by the Innovative Genomics Institute, the Allen Foundation, and the Chan Zuckerberg Biohub.

## Author Contributions

C.H. and J.F.B. designed the study. C.H., R.K., and I.F. collected samples, extracted DNA, and performed manual binning. C.H. performed automated binning, bin selection, bin curation and dereplication, phylogenetic analysis, abundance and community comparison analysis, metabolic analysis, and cell count distribution analysis. M.W. prepared cryo-TEM grids and collected cryo-TEM data. R.K. performed ordination analysis. C.H. and J.F.B. drafted the manuscript. All authors reviewed the manuscript.

## Supplementary Information

**SI Fig 1:**
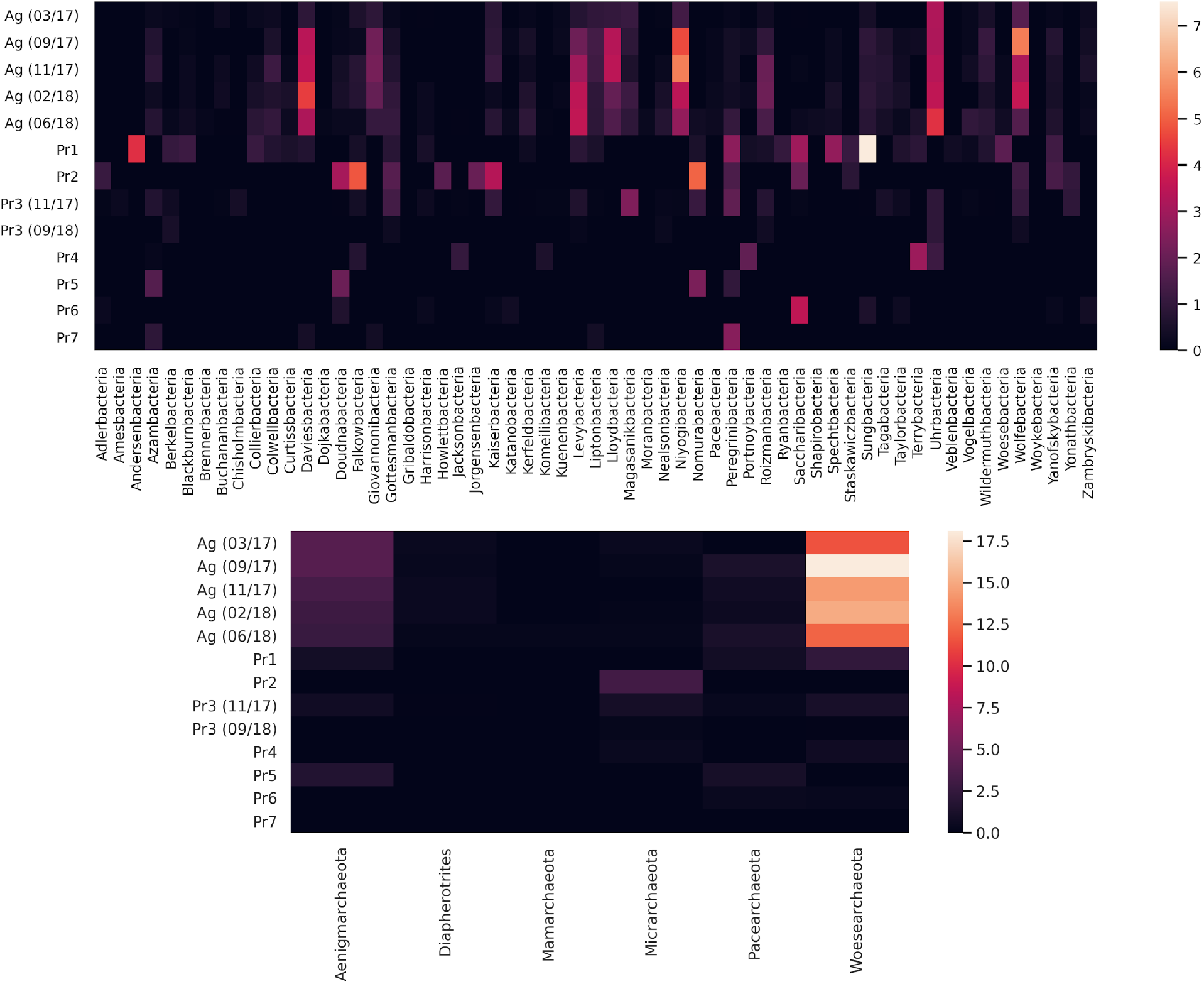
Distribution of CPR (top) and DPANN (bottom) phylum-level lineages across groundwater sites. Color/legend indicate relative coverage values (percentages).

**SI Fig 2:**
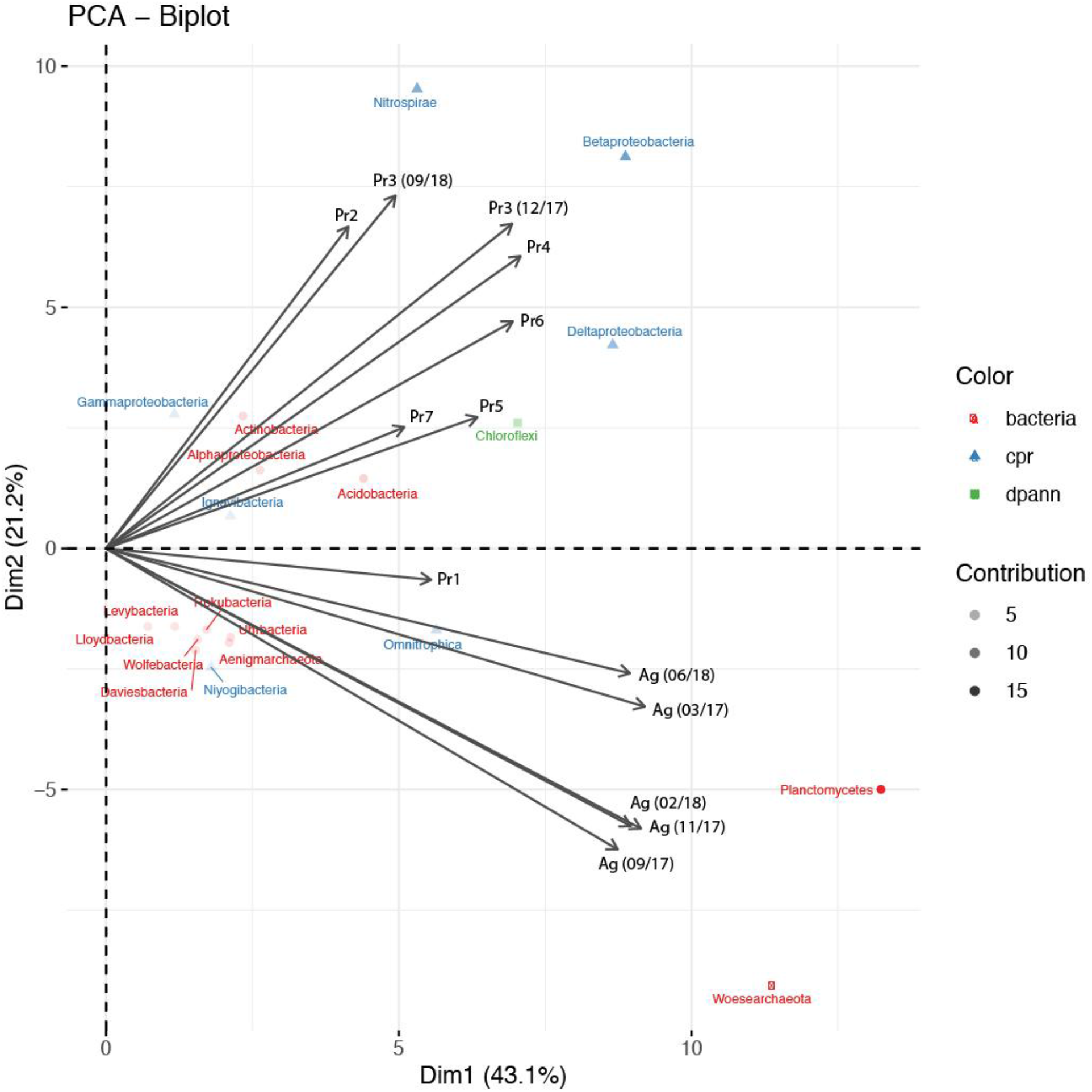
A biplot representation of phylum-level lineages in ordination space, with arrows showing the direction of greatest gradient change according to site. The transparency of the points reflects the contribution of the phylum to the principal components. *Ag* clusters separately from all pristine sites based upon strong and unique representation of organisms from Planctomycetes, Woesearchaeota, and several CPR lineages. *Pr1* is the pristine site that clusters closest to *Ag* based due to shared representation of Omnitrophica, while the remaining six pristine sites are differentiated by representation of Beta- and Deltaproteobacteria, Chloroflexi, and Nitrospirae.

**SI Fig 3:**
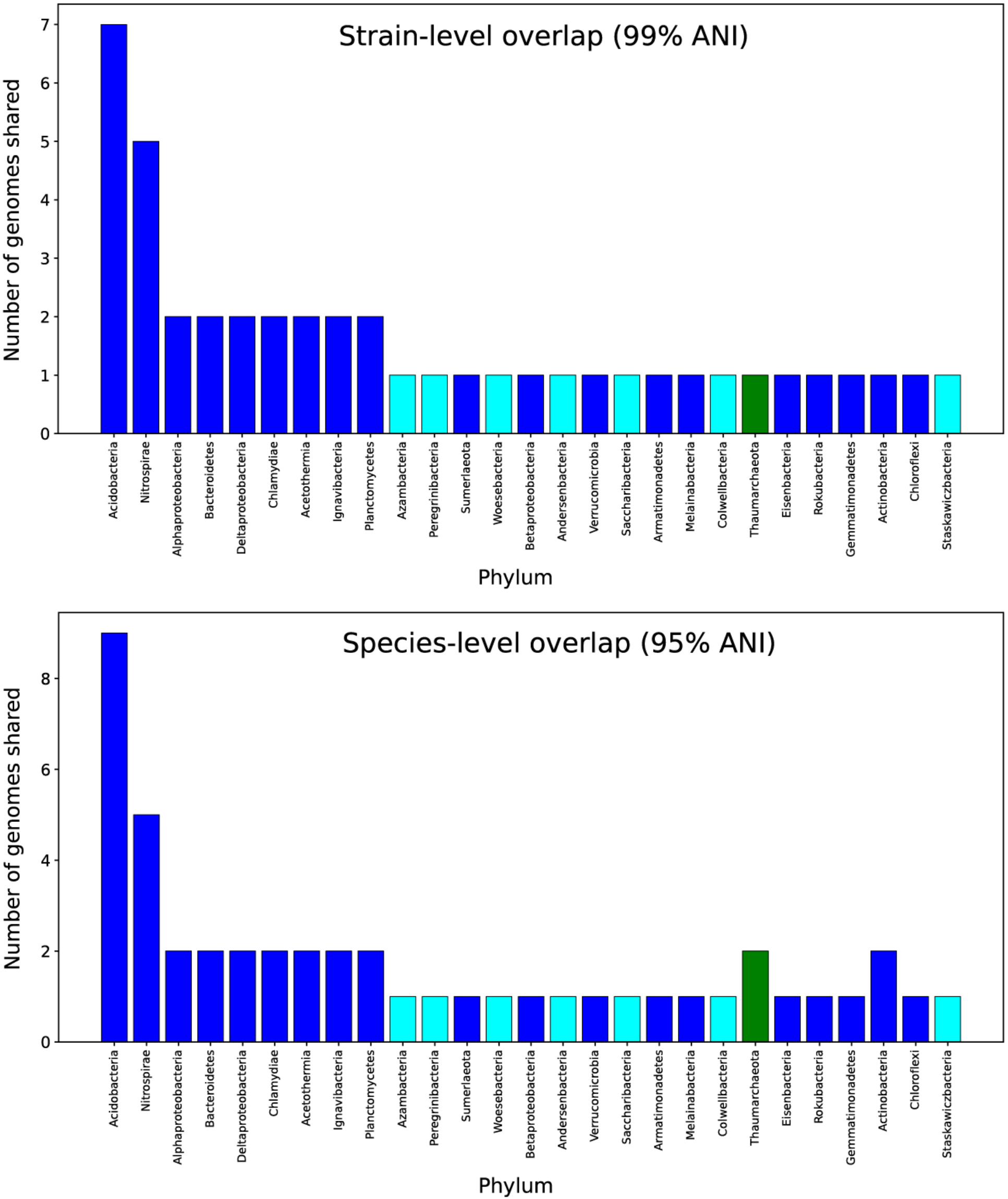
Genome similarity at the strain (>99% ANI) and species (>95% ANI) level between *Pr1* and *Pr7* genomes. Blue bars indicate non-CPR bacteria, aqua bars indicate CPR bacteria, and green bars indicate archaea.

**SI Fig 4:**
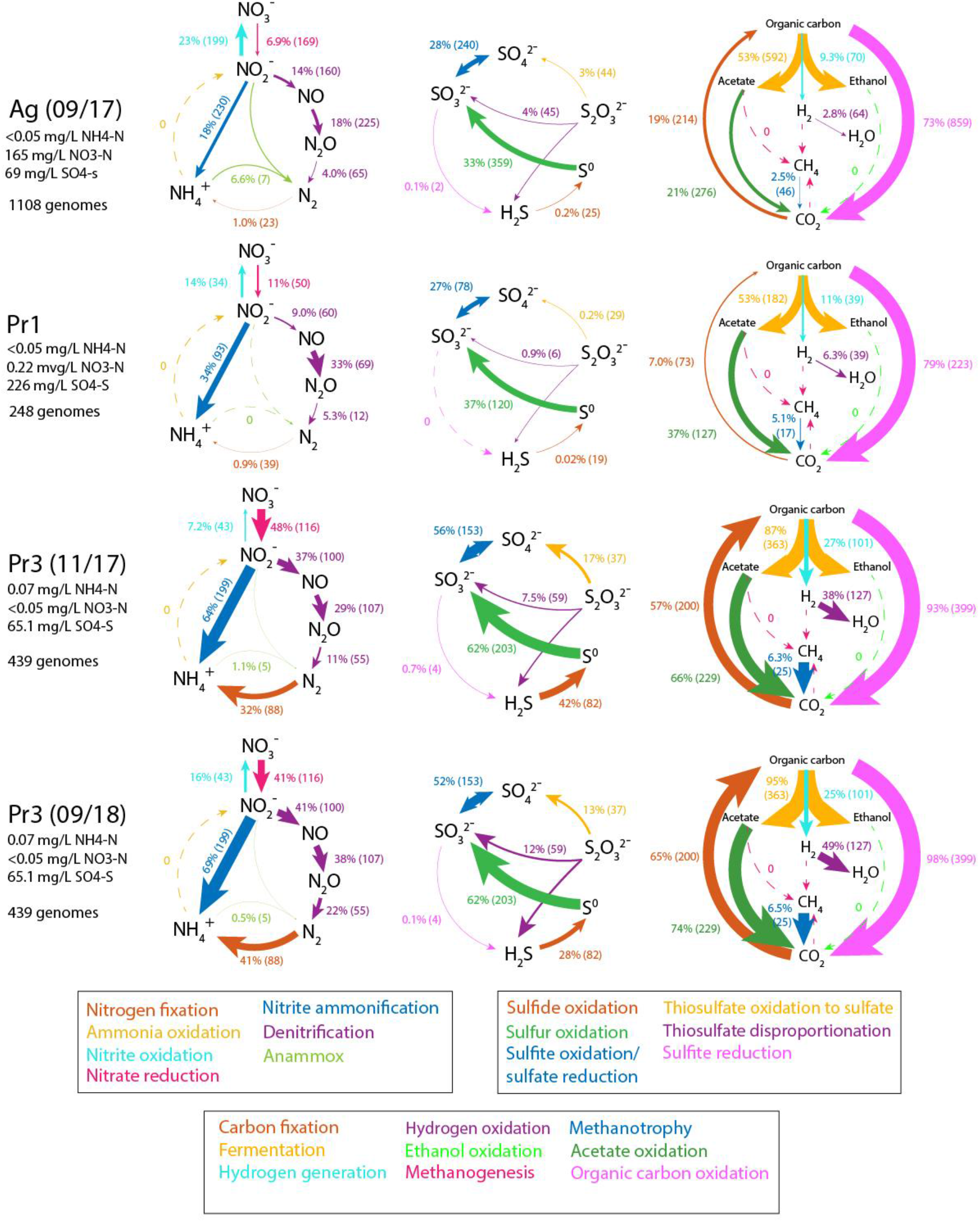
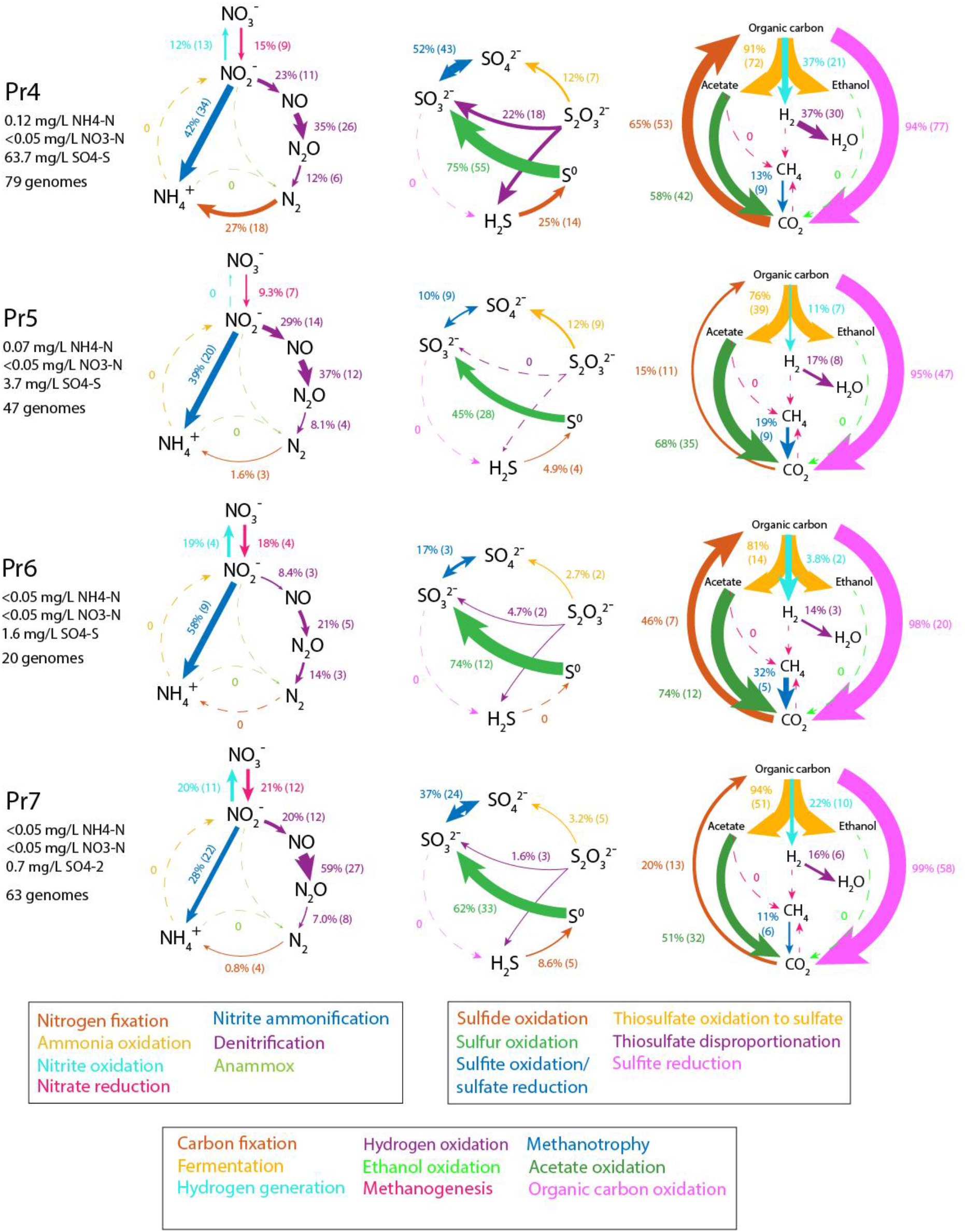
Community-level cycling of nitrogen, sulfur, and carbon in the eight groundwater communities sampled in this study. Listed next to each metabolic step are the total relative abundance of all genomes capable of carrying out the step, and the number of genomes containing the capacity for that step. Arrow sizes are drawn proportional to the total relative abundance of genomes capable of carrying out the metabolic step.

**SI Fig 5:**
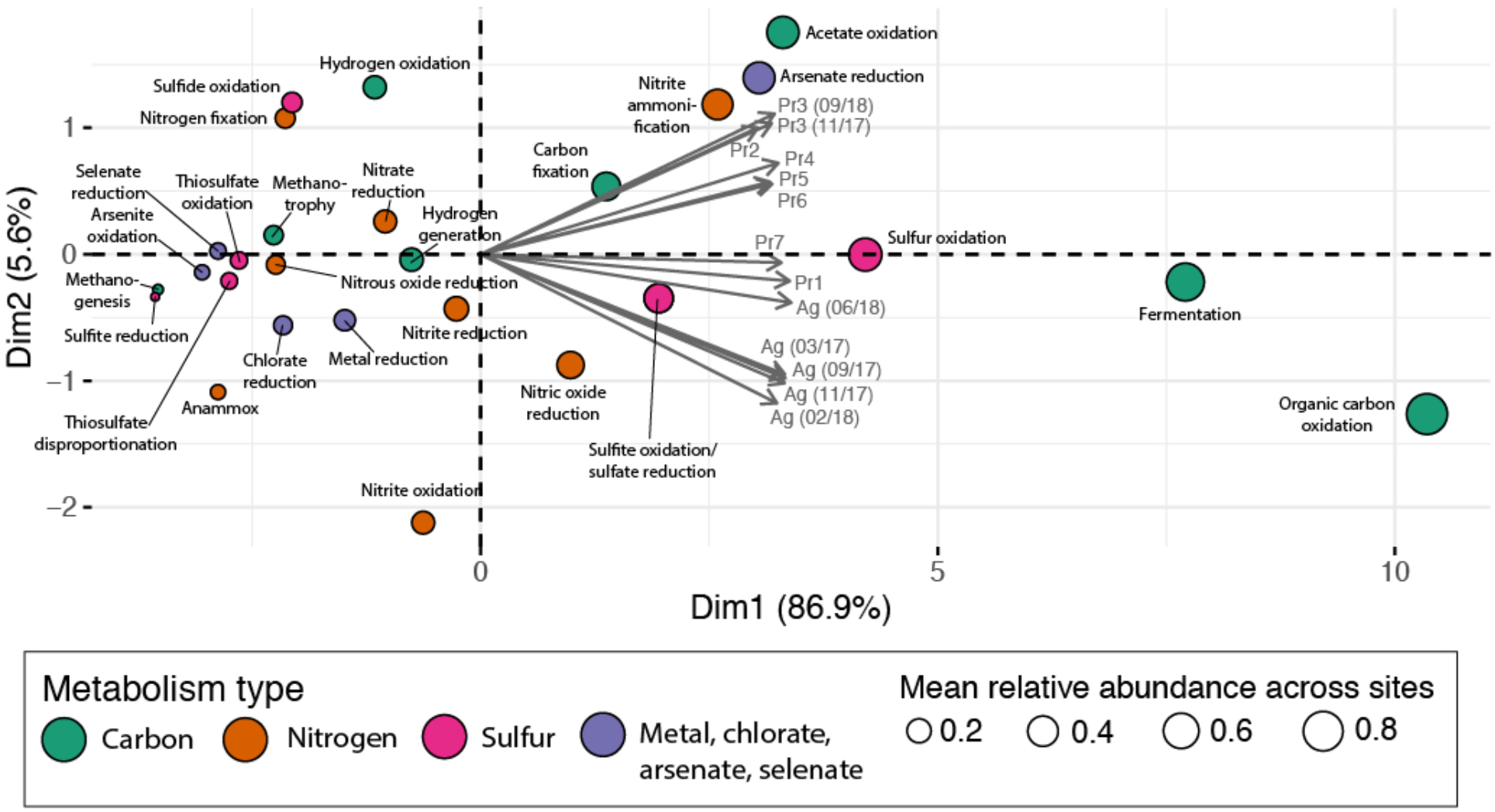
Depiction of total relative coverages of different metabolic capacities, in principal component space. Principal component analysis was performed on the relative abundance of organisms with specific metabolic capacities across all sites. Organic carbon oxidation (>70% abundance in all sites) and fermentation (>50% abundance in all sites) account for the majority of variance in the data, while the divide between *Pr7, Pr1*, and *Ag* and the other sites is determined by a range of factors including lower levels of acetate oxidation and carbon fixation than the other sites.

**SI Fig 6:**
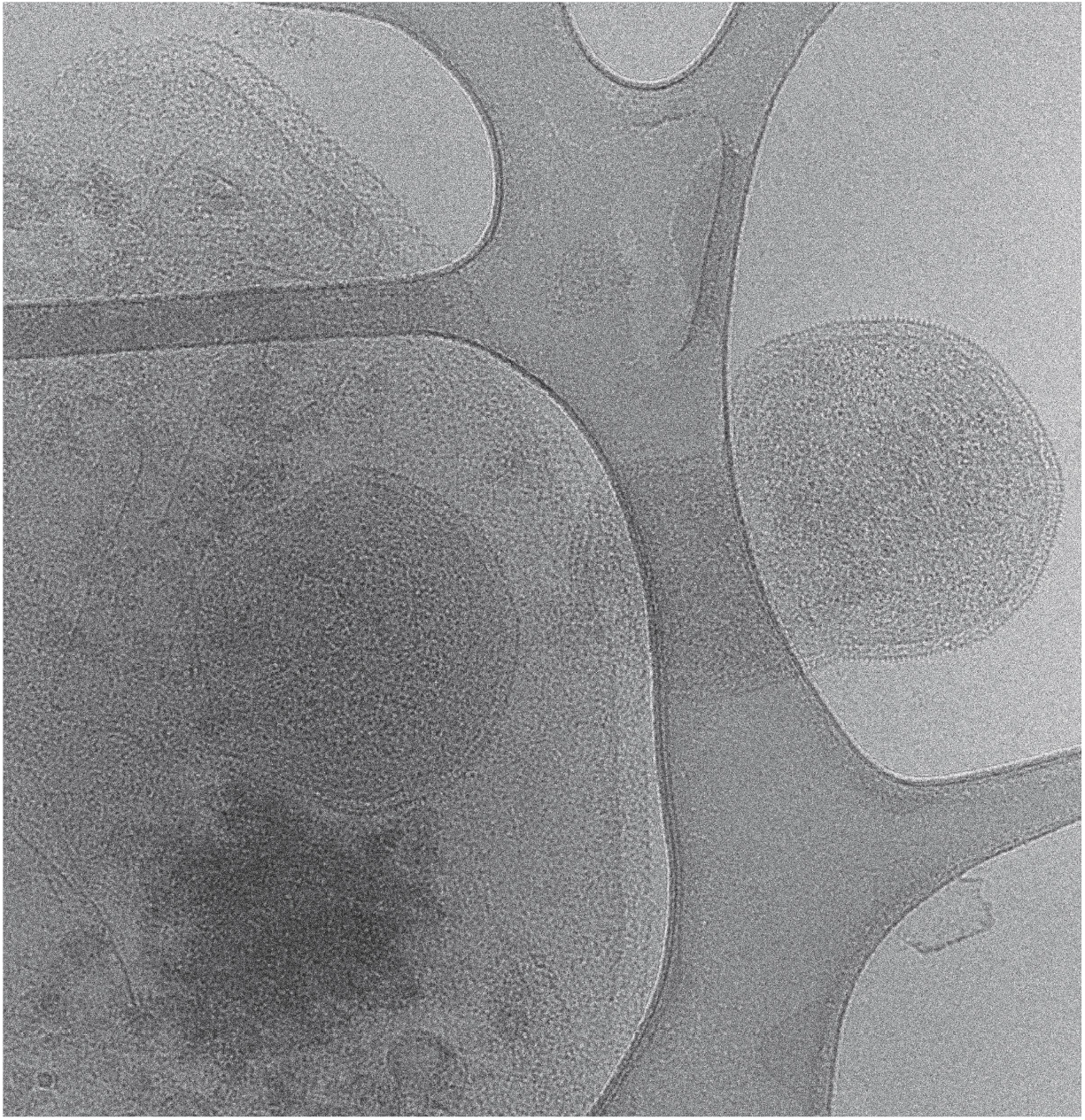
Cryo-TEM image of ultra-small cells associated with dead/lysed cells in *Ag* groundwater concentrated by tangential flow filtration.

**SI Fig 7:**
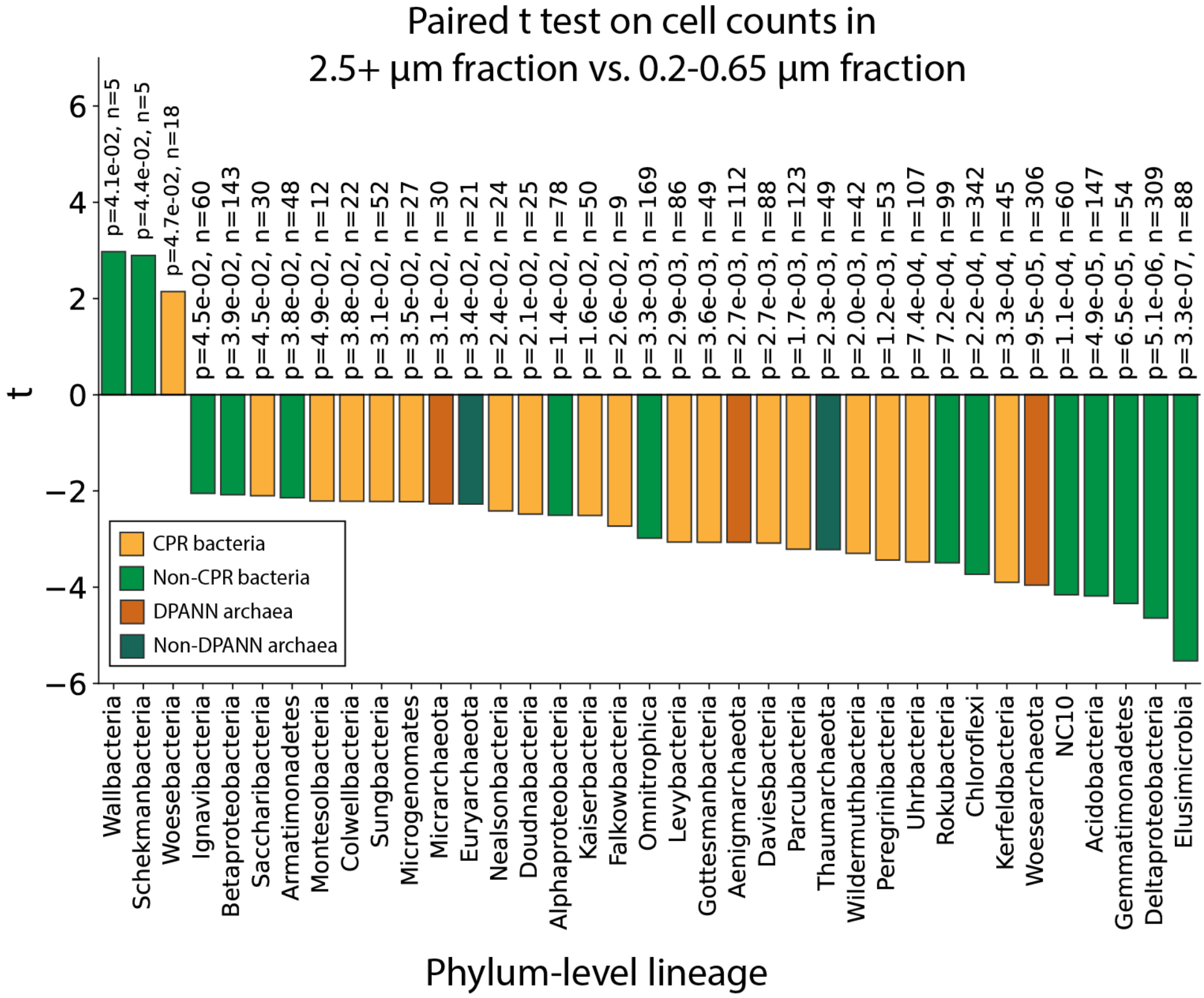
Results from a paired t-test on estimated cell counts of genomes in the 2.5+ μm versus the 0.2-0.65 μm size fractions after serial size filtration. A positive t statistic indicates enrichment on the 2.5+ μm compared to the 0.2-0.65 μm size fraction. Values listed by each bar are the calculated p value (top value) and sample size (bottom value) for each phylum-level lineage. Notably, CPR bacterial lineage Woesebacteria stands out as being enriched in the 2.5+ μm compared to the 0.2-0.65 μm size fraction.

## Supplementary Notes

### Supplementary Note 1: Differences in *Pr3* community composition between time points

From Nov 2017 to Sept 2018, *Pr3* groundwater showed a drop from ~17% to ~3% CPR abundance, 3.5% to 0.35% DPANN abundance, and 3.3% to 0.5% abundance of non-DPANN archaea, offset by an increase in non-CPR bacteria (Fig 2A and 1C). Several CPR and DPANN phylum-level lineages detected in Nov 2017 were no longer detectable in Sept 2018 (Fig 2A). Additionally, we observe a shift from Deltaproteobacteria and Betaproteobacteria as the most abundant organisms to Nitrospirae (Fig 1C). This observed change likely relates to sampling prior to versus after onset of the rainy season, and consequential changes in groundwater chemistry.

### Supplementary Note 2: Potential index hopping between Pr1 and Pr7

*Pr1* and *Pr7* was the only pair of analyzed sites that shared more than a few strains (44 pairs of genomes with >99% ANI). These two sites are separated by Putah Creek, a major hydrological feature, and so are unlikely to be fed from the same aquifer. Although DNA from *Pr1* and *Pr7* was sequenced on the same NovaSeq 6000 lane, we do not attribute this overlap to index hopping, as dual indexing was used and reads with mismatched indices were not analyzed, reducing the already low incidence of index hopping (<2% of reads) significantly. Furthermore, though the 44 genome pairs share >99% ANI, they are not identical, differing in sequence by up to 10,000 bp per Mbp of genome.

## Supplementary Methods

### Ordination analysis

Principal component analysis (PCA) was performed on rpS3 relative coverage values and on metabolic capacities of whole communities (the summed relative coverage values of genomes encoding a particular metabolic transformation). PCA was performed using the FactoMineR package^1^ and visualized with factoextra^2^. Relative abundance values were scaled to unit variance prior to the calculation of the principal components. Non-metric multidimensional scaling (NMDS) analysis was performed on normalized read counts (reads per million total reads) for all genomes from *Ag* groundwater, based on read mapping with BBMap (Bushnell B. – sourceforge.net/projects/bbmap/). NMDS analysis was performed using the metaMDS function in the Vegan package for R^3^, with default parameters. Briefly, the data was transformed using Wisconsin double standardization of the square root of the matrix, followed by construction of Bray-Curtis dissimilarity matrix, then an nMDS with 20 random starts. Finally, the results were scaled to maximize variation to the first principal component. Results were visualized using the ggplot2 package for R^4^.

### Phylogenetic classification

Genomes with a clear taxonomic classification based on the internal ggKbase database (>50% of the genome sequence had a clear scaffold-level taxonomic winner, based on best matches of protein sequences to those in genomes of a taxonomically comprehensive databases) were classified according to their predicted ggKbase taxonomy. For genomes without a clear predicted ggKbase taxonomy, phylogenetic analysis was carried out using several marker sets: concatenated ribosomal proteins (encoded by a syntenic block of genes and selected to avoid binning error chimeras), rpS3 proteins, and 16S rRNA genes (for CPR bacteria). Reference sequences for all phylogenetic trees were taken from previous studies that recovered many high quality CPR and DPANN genomes^5–8^.

The concatenated ribosomal protein set for bacteria includes 15 proteins (L2, L3, L4, L5, L6, L14, L15, L18, L22, L24, S3, S8, S10, S17, and S19) while the archaeal set includes 14 proteins (the bacterial set without S10, which is missing from many archaeal genomes). Ribosomal proteins were identified by searching predicted ORFs against ribosomal protein databases usearch^9^. Each ribosomal protein was aligned to its Pfam HMM model using hmmalign from HMMer^10^ (version 3.3), alignments were converted from Stockholm format to FASTA, and alignment insert columns were removed. All ribosomal proteins were concatenated together, and concatenated sequences with an ungapped length of over 1,100 amino acid residues were combined with reference sequences to build a maximum likelihood tree using IQ-Tree (version 1.6.12) (iqtree -s <alignmentfile> -st AA -nt 48 -bb 1000 -m LG+G4+FO+I).

For rpS3 gene phylogenetic analysis, rpS3 genes were identified using a custom HMM with an HMM alignment score cutoff of 40^11^. Identified rpS3 genes were aligned with rpS3 reference sequences using mafft^12^ (default parameters) and columns with >95% gaps were removed with trimal^13^. The alignment was used to build a maximum likelihood tree using IQ-Tree (iqtree -s <alignmentfile> -st AA -nt 48 -bb 1000 -m LG+G4+FO+I).

For 16S rRNA gene phylogenetic analysis of CPR bacterial genomes, 16S rRNA genes were identified using a custom HMM^5^ and insertions of 10 bp or greater were removed. Sequences with lengths of >800bp were used for phylogenetic analysis. SSU-align was used to align 16S sequences from this study with reference sequences from the previous studies mentioned above as well as CPR bacteria sequences from SILVA database^14^. The resulting alignment was used to build a maximum likelihood tree with RAxML-HPC BlackBox^15^ (8.2.12) on the CIPRES Science Gateway^16^ with an LG protein substitution matrix.

Genomes forming novel phylum-level lineages Genascibacteria and Montesolbacteria were identified based on the following criteria: (1) they formed a monophyletic group in the 16S rRNA gene phylogeny, (2) 16S rRNA genes were at least 24% divergent from the closest representatives, (3) they were also supported by the concatenated ribosomal protein phylogeny, and (4) more than one representative draft genome was available. Genascibacteria and Montesolbacteria genomes and ANI with closest 16S rRNA hits from SILVA^14^ are listed in SI Table 3.

